# Lysine methyltransferase SET7 links cardiometabolic risk to endothelial dysfunction by dysregulating mRNA splicing and eNOS-CaM interaction

**DOI:** 10.64898/2026.08.02.742358

**Authors:** Julia Sánchez-Ceinos, Georgios Filis, Jingyi Zhang, Magnus E. Jakobsson, Ákos Végvári, Cheukyau Luk, Emelie Carlestål, Carolina E. Hagberg, Oskar Kövamees, Francesco Cosentino

## Abstract

**BACKGROUND:** Lysine methyltransferase SET7 activates gene expression through mono- methylation of histone H3 at lysine 4 (H3K4me1) and modulates protein function via mono- methylation of non-histone proteins (Kme1). However, its role and molecular targets in endothelial dysfunction associated with cardiometabolic disorders remain unknown.

**METHODS:** Endothelial-specific *Setd7* knockout (*Setd7*EC-KO) mice were generated and endothelial function assessed in WT and *Setd7*EC-KO mice after high-fat diet (HFD). Human aortic endothelial cells (HAECs) were used to investigate SET7 expression and function under metabolic stress. SET7-dependent histone and non-histone targets were identified by proteomic and ChIP analyses. Insights from these datasets guided the design of bioinformatic, molecular, and functional studies to define their regulatory mechanisms. Clinical relevance was evaluated in human arteries.

**RESULTS:** HFD selectively increased endothelial SET7 expression in WT mouse aortas. Despite comparable metabolic abnormalities, *Setd7*EC-KO mice were protected from HFD- induced endothelial dysfunction, oxidative stress, and inflammation. In HAECs, high glucose emerged as the strongest inducer of SET7 expression, promoting pro-inflammatory and pro- oxidant gene expression, monocyte adhesion, and ROS generation. These effects were reproduced by overexpression of catalytically active SET7 and reversed by its inhibition or silencing. Proteomic and ChIP analyses revealed that SET7-dependent H3K4me1 activates transcription of spliceosome components, linking aberrant mRNA splicing to endothelial inflammation and oxidative stress. Moreover, Kme1-proteomics identified endothelial nitric oxide synthase (eNOS) as a direct SET7 substrate. Bioinformatic analyses and mutagenesis experiments demonstrated that SET7-mediated mono-methylation of eNOS at K494 disrupts calmodulin (CaM) binding and impairs NO synthesis. These molecular signatures were also observed in internal mammary arteries from patients with vascular disease and hyperglycemia.

**CONCLUSIONS:** SET7 drives endothelial dysfunction through a dual mechanism: 1) H3K4me1-dependent activation of splicing machinery triggering inflammation and oxidative stress, and 2) eNOS mono-methylation at K494 reducing NO bioavailability. Targeting SET7 may therefore represent a promising avenue to safeguard endothelial homeostasis in cardiometabolic disease.

**GRAPHIC ABSTRACT:** A graphic abstract is available for this article.

## Introduction

Atherosclerotic cardiovascular disease (ASCVD) remains the leading cause of morbidity and mortality worldwide, and its prevalence is projected to rise substantially in the coming decades^1^. Metabolic disorders such as obesity and type 2 diabetes (T2D) represent major global health challenges that substantially contribute to the growing burden of ASCVD^2^. Strategies that preserve vascular health by targeting early alterations in the vasculature are therefore essential for preventing or delaying CV complications in these high-risk populations. Endothelial dysfunction, characterised by impaired vasoreactivity, reduced nitric oxide (NO) bioavailability, inflammation, and oxidative stress, plays a central role in the initiation and development of ASCVD^3^. However, the precise molecular mechanisms triggering the activation of these pro-atherogenic conditions in cardiometabolic disease are still incompletely understood.

The SET domain-containing lysine methyltransferase 7 (SET7), also known as SET9^4^, SET7/9^5^, or KMT7^6^, has traditionally been recognised for catalysing mono-methylation of histone H3 at lysine 4 (H3K4me1) in gene promoter regions. Such epigenetic modification leads to chromatin accessibility and transcriptional activation^7–9^. More recently, accumulating evidence indicated that SET7 also targets a wide range of non-histone proteins^10,11^, including p53^12,13^, NF-kB p65^14,15^, STAT3^16^, YAP^17,18^, DNMT1^19,20^, FOXO3^21,22^, SIRT1^23^, and SUV39H1^24^. Mono-methylation of lysine residues (Kme1) on non-histone proteins can influence protein interactions and stability, subcellular localisation, enzymatic activity, and crosstalk with other post-translational modifications^25,26^, extending the regulatory action of SET7 as a versatile modulator of protein function far beyond chromatin remodelling ^27^. Through these combined histone and non-histone activities, SET7 has been implicated in diverse biological processes, including inflammation^28–30^ and oxidative stress^31–33^, across multiple cell types and disease contexts. Nevertheless, the role of SET7 and its specific histone (H3K4me1) and non-histone (Kme1) targets in endothelial dysfunction associated with cardiometabolic risk remains to be fully elucidated.

Here, we address these open questions by integrating chromatin, transcriptomic, proteomic, and functional approaches across an endothelial-specific *Setd7* knockout (*Setd7*EC-KO) mouse model subjected to high-fat diet (HFD), human aortic endothelial cells (HAECs) exposed to metabolic stressors that mimic HFD and modulation of SET7 expression, as well as in arteries from patients with ASCVD and hyperglycemia. All these analyses consistently identify SET7 as a pivotal mediator of endothelial dysfunction under metabolic impairment. Indeed, we demonstrate that HFD and elevated glucose levels induce SET7 expression in aortic endothelial cells, which in turn promotes endothelial damage via two complementary pathways: 1) H3K4me1-dependent transcriptional activation of spliceosome components that contributes to inflammatory and oxidative phenotypes; and 2) direct mono-methylation of endothelial nitric oxide synthase (eNOS) at lysine 494 that disrupting calmodulin (CaM) binding reduces NO production. Interestingly, both mechanisms are also present in arteries from patients with ASCVD and hyperglycemia. Taken together our results define previously unrecognised molecular pathways linking metabolic imbalance to vascular injury and establish SET7 as a promising therapeutic candidate for preventing or mitigating cardiometabolic vascular complications.

## METHODS

### Data Availability

A full description of all experimental procedures and materials used in this study is provided in the **Supplemental Methods** section. Information on key experimental materials is listed in the Major Resources Table in the **Supplemental Material**. All data generated and/or analysed during this study are included in this article (and its **Supplemental Material**). Further information about the data and materials used is available from the corresponding authors upon reasonable request. All mass spectrometry proteomics data generated in this study have been deposited in the ProteomeXchange Consortium via the PRIDE repository34,35, with the dataset identifier PXD074145.

### Animal Model

Endothelial-specific, tamoxifen-inducible *Setd7* knockout mice (*Setd7*EC-KO) were generated by crossing *Setd7fl/fl* mice with Tek-Cre mice. *Setd7fl/fl* littermates lacking Cre served as wild-type (WT) controls. Mice received tamoxifen (40 mg/kg) for 5 weeks to induce recombination and were then fed a standard (STD, 10% kcal fat) or high-fat diet (HFD, 60% kcal fat) for 12 weeks. Aortas and blood were collected at 12–18 weeks of age. Genotypes were verified by conventional PCR (**Figure S1A**), and efficient and selective deletion of SET7 in the endothelium was validated by RT-PCR and immunofluorescence (**Figures S1B** and **S1C**). All procedures were approved by the Karolinska Institutet Animal Care and Use Committee. Blood glucose was measured immediately after collection. Plasma insulin, lipids, and free fatty acids (FFAs) were quantified using commercial assays. Insulin resistance was calculated using HOMA-IR

### Wire Myograph

Aortic rings were mounted in a wire myograph system and equilibrated under physiological tension^36^. Endothelium-dependent and -independent relaxations were assessed using acetylcholine (Ach, 10–9–10–4.5 M) and sodium nitroprusside (SNP, 10–10–10–5.5 M), respectively, after phenylephrine-induced contraction (PE, 10–6 M). Reactive oxygen species (ROS) scavenger experiments were performed in vessels from WT-HFD mice.

### Measurement of ROS

Superoxide (O2−) and peroxynitrite (ONOO^-^) in mouse aortas were quantified by electron spin resonance (ESR) spectroscopy using specific spin probes^36^. Intracellular ROS in HAECs were measured using 2’-7’-dichlorodihydrofluorescein diacetate (DCFH-DA) fluorescence and normalised to protein content.

### Immunohistochemistry of Human and Mouse Arteries

Human internal mammary arteries and mouse aortic rings were fixed, paraffin-embedded, sectioned, and subjected to antigen retrieval^36^. Sections were incubated with primary antibodies and visualised using fluorophore-conjugated secondary antibodies (**Supplemental Material**). Nuclei were counterstained with Hoechst.

### Cell Culture and Experimental Treatments

Human aortic endothelial cells (HAECs) were cultured in endothelial growth medium under standard conditions^36^ and used between passages 1–8. Mycoplasma testing was performed routinely. HAECs were exposed to high glucose (HG; 25 mM), insulin (100 nM), fatty acids (oleate or palmitate, 250 µM), or LDL cholesterol (20 nM) for 24 hours to mimic HFD conditions^36,37^. SET7 activity was inhibited pharmacologically using cyproheptadine^38^ (10 µM) or by siRNA, and mRNA splicing was inhibited using pladienolide B^37^ (PladB, 100 pM). Cyproheptadine and PladB were selected at the highest concentration that did not compromise cell viability (**Figures S2A** and **B**, respectively). WT and mutant (MT) SET7 and eNOS constructs were introduced using plasmid transfection^37^. Expression efficiency was confirmed by RT-PCR and/or immunoblotting.

### RNA Extraction and RT-PCR

Total RNA was extracted, reverse transcribed, and analysed by SYBR Green RT-PCR using previously validated primers (**Table S1**). Expression was normalised to housekeeping genes using the 2-ΔCt method^36,37,39,40^.

### Monocyte Adhesion Assay

THP-1 human monocytes were labelled with calcein in green and incubated with HAECs. Adhesion was quantified by confocal microscopy.

### Immunoprecipitation (IP) of Kme1-modified Proteins

Kme1-modified proteins were immunoprecipitated using a specific antibody and magnetic beads, followed by downstream proteomic analysis. The anti-Kme1 antibody was in-house manufactured, validated by ELISA and dot blot analyses (**Figures S3A** and **S3B**), and kindly provided by Magnus E. Jakobsson (Malmö University, Sweden). Dot blot analyses of samples collected at different steps of the IP process, together with recombinant methylated BSA as a positive control, were performed to verify the efficiency of the IP (**Figure S3C**).

### Liquid Chromatography-Tandem Mass Spectrometry (LC-MS/MS) Proteomic Analysis

Proteins in the input and IP fractions were digested with trypsin and analysed by LC-MS/MS. Data were processed using Proteome Discoverer with label-free quantification and False Discovery Rate filtering.

### Chromatin immunoprecipitation coupled with quantitative PCR (ChIP-qPCR)

Chromatin immunoprecipitation was performed as described before^36^ using antibodies against H3K4me1, followed by qPCR analysis of promoter regions of target genes using specific primers (**Table S1**).

### Site-directed Mutagenesis of eNOS

Construct encoding eNOS with lysine-to-isoleucine substitution at residue 494 (eNOS-K494I) was generated by site-directed mutagenesis and verified by Sanger sequencing.

### Bioinformatics Analyses

Pathway enrichment was performed using Reactome^41^ and protein interaction networks using STRING^42^. Methylation sites were predicted using MethylSight^43^. Protein structures were retrieved from RCSB Protein Data Bank^44^ or AlphaFold^45^. Docking was performed using ClusPro^46^ and visualised in ChimeraX^47^. Structural effects of mutations were assessed by root-mean-squared analysis.

### Patient Recruitment and Collection of Internal Mammary Arteries (IMAs)

Internal mammary arteries (IMAs) were obtained from patients undergoing bypass surgery (n=12) and stratified by glycemic status (**Table 1**). Tissues were processed for histological analysis.

**Table 1.**
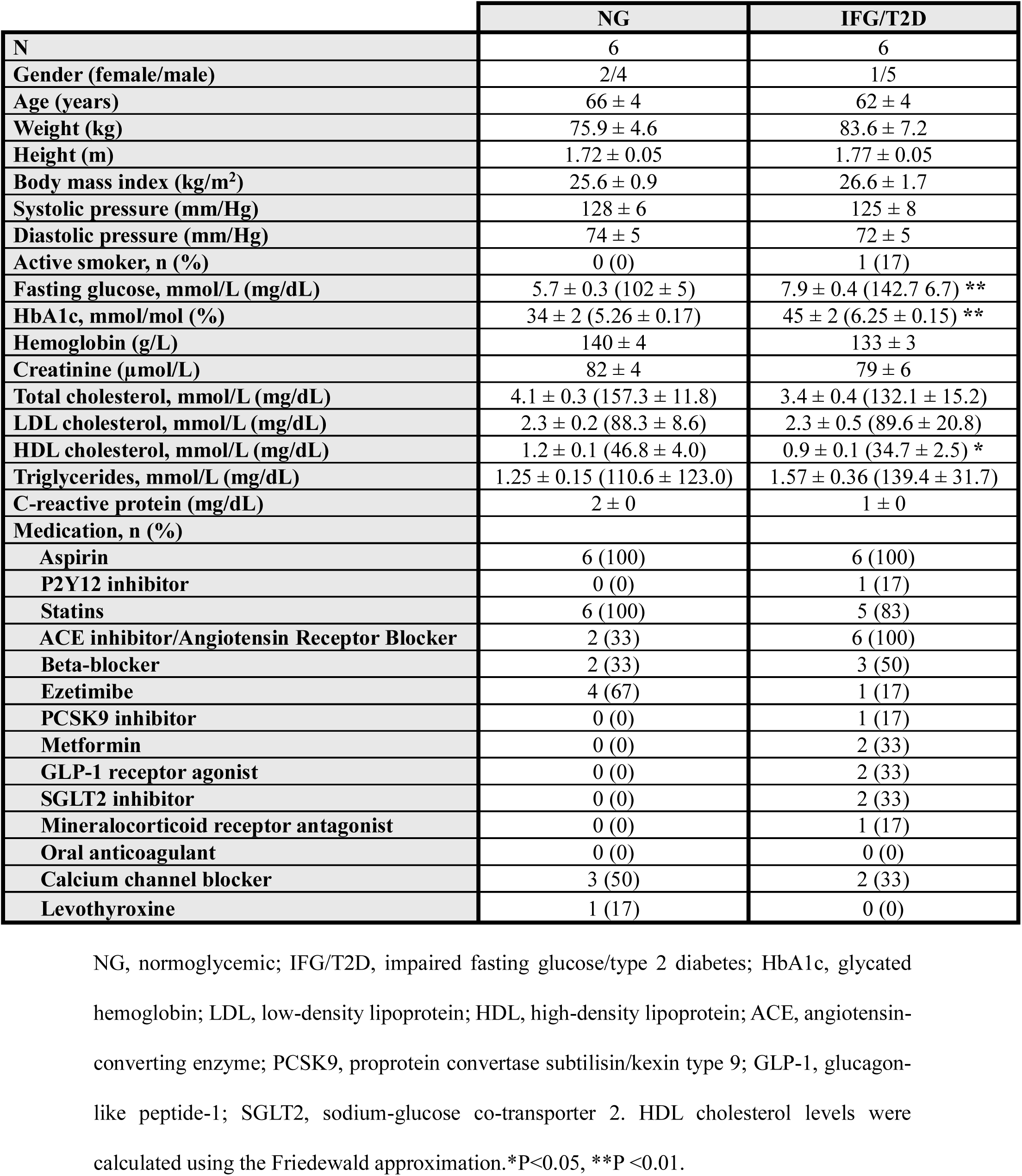
Anthropometric and biochemical characteristics of study subjects.

|  | NG | IFG/T2D |
| --- | --- | --- |
| **N** | 6 | 6 |
| **Gender (female/male)** | 2/4 | 1/5 |
| **Age (years)** | 66 ± 4 | 62 ± 4 |
| **Weight (kg)** | 75.9 ± 4.6 | 83.6 ± 7.2 |
| **Height (m)** | 1.72 ± 0.05 | 1.77 ± 0.05 |
| **Body mass index (kg/m<sup>2</sup>)** | 25.6 ± 0.9 | 26.6 ± 1.7 |
| **Systolic pressure (mm/Hg)** | 128 ± 6 | 125 ± 8 |
| **Diastolic pressure (mm/Hg)** | 74 ± 5 | 72 ± 5 |
| **Active smoker, n (%)** | 0 (0) | 1 (17) |
| **Fasting glucose, mmol/L (mg/dL)** | 5.7 ± 0.3 (102 ± 5) | 7.9 ± 0.4 (142.7 6.7) ** |
| **HbA1c, mmol/mol (%)** | 34 ± 2 (5.26 ± 0.17) | 45 ± 2 (6.25 ± 0.15) ** |
| **Hemoglobin (g/L)** | 140 ± 4 | 133 ± 3 |
| **Creatinine (μmol/L)** | 82 ± 4 | 79 ± 6 |
| **Total cholesterol, mmol/L (mg/dL)** | 4.1 ± 0.3 (157.3 ± 11.8) | 3.4 ± 0.4 (132.1 ± 15.2) |
| **LDL cholesterol, mmol/L (mg/dL)** | 2.3 ± 0.2 (88.3 ± 8.6) | 2.3 ± 0.5 (89.6 ± 20.8) |
| **HDL cholesterol, mmol/L (mg/dL)** | 1.2 ± 0.1 (46.8 ± 4.0) | 0.9 ± 0.1 (34.7 ± 2.5) * |
| **Triglycerides, mmol/L (mg/dL)** | 1.25 ± 0.15 (110.6 ± 123.0) | 1.57 ± 0.36 (139.4 ± 31.7) |
| **C-reactive protein (mg/dL)** | 2 ± 0 | 1 ± 0 |
| **Medication, n (%)** |  |  |
| **Aspirin** | 6 (100) | 6 (100) |
| **P2Y12 inhibitor** | 0 (0) | 1 (17) |
| **Statins** | 6 (100) | 5 (83) |
| **ACE inhibitor/Angiotensin Receptor Blocker** | 2 (33) | 6 (100) |
| **Beta-blocker** | 2 (33) | 3 (50) |
| **Ezetimibe** | 4 (67) | 1 (17) |
| **PCSK9 inhibitor** | 0 (0) | 1 (17) |
| **Metformin** | 0 (0) | 2 (33) |
| **GLP-1 receptor agonist** | 0 (0) | 2 (33) |
| **SGLT2 inhibitor** | 0 (0) | 2 (33) |
| **Mineralocorticoid receptor antagonist** | 0 (0) | 1 (17) |
| **Oral anticoagulant** | 0 (0) | 0 (0) |
| **Calcium channel blocker** | 3 (50) | 2 (33) |
| **Levothyroxine** | 1 (17) | 0 (0) |

NG, normoglycemic; IFG/T2D, impaired fasting glucose/type 2 diabetes; HbA1c, glycated
hemoglobin; LDL, low-density lipoprotein; HDL, high-density lipoprotein; ACE, angiotensin-
converting enzyme; PCSK9, proprotein convertase subtilisin/kexin type 9; GLP-1, glucagon-
like peptide-1; SGLT2, sodium-glucose co-transporter 2. HDL cholesterol levels were
calculated using the Friedewald approximation.\*P<0.05, \*\*P<0.01.

### Statistical Analysis

Data are presented as mean ± SEM. Statistical significance was assessed using t-tests, ANOVA, or non-parametric equivalents as appropriate. Correlations were evaluated using Pearson’s or Spearmańs correlation analyses for parametric and non-parametric data, respectively. P < 0.05 was considered significant.

## RESULTS

### Lysine Methyltransferase SET7 is Upregulated in the Endothelium of Mice Exposed to High-Fat Diet

To investigate the role of lysine methyltransferase SET7 in endothelial dysfunction associated with metabolic disturbance, WT C57BL/6 mice were fed either standard diet (STD) or high-fat diet (HFD) for 12 weeks (**Figure 1**). Then, mice were sacrificed and their aortas collected for endothelial cell isolation and confocal microscopy studies (**Figure 1A**). RT-PCR analyses showed a significant upregulation of *Setd7* (encoding SET7) expression in endothelial cells from HFD animals compared with their STD littermates (**Figure 1B**, left panel). In contrast, *Setd7* mRNA levels remained unchanged in cells from the media and adventitia layers of the same aortas (**Figure 1B**, right panel). This endothelium-specific transcriptional upregulation was confirmed at the protein level by immunofluorescence co-staining for SET7 and the endothelial marker CD31 in aortic tissue sections (**Figure 1C**).

**Figure 1.**
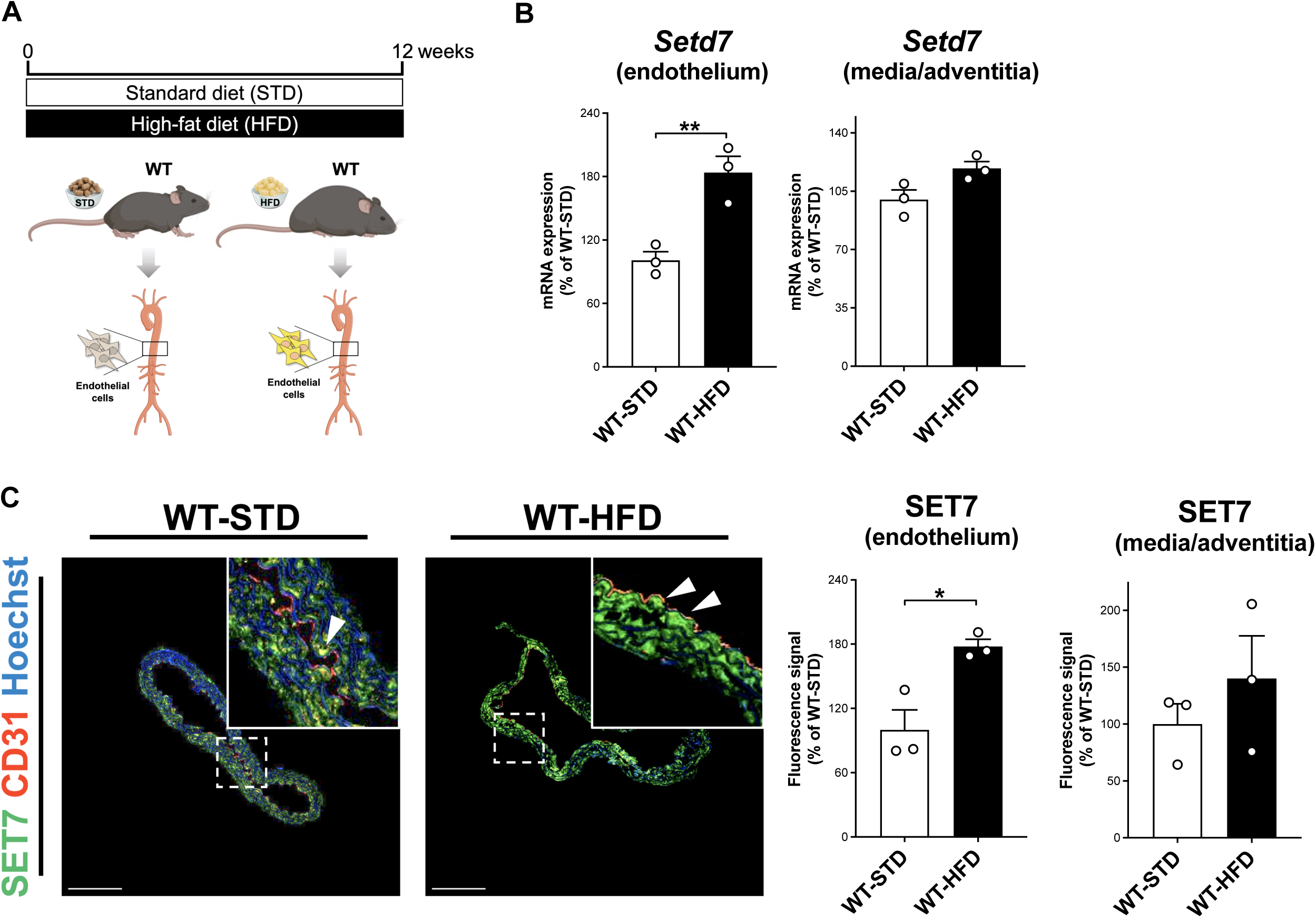
Lysine methyltransferase SET7 is upregulated in the endothelium of mice exposed to high-fat diet. **A**, Schematic representation of the experimental animal design. **B**, mRNA levels of *Setd7*, and **C**, representative confocal images and quantification of SET7 (green) in the endothelium and the media/adventitia layers of aortas from WT mice fed with standard diet (STD) or a high-fat diet (HFD). The endothelium was visualized by CD31 immunostaining (red) and nuclei were counterstained with Hoechst (blue). Scale bar = 200 μm. *P<0.05, **P<0.01.

### Endothelium-Specific Deletion of *Setd7* Preserves Endothelial Function by Attenuating Oxidative Stress and Inflammation

Based on these findings, to elucidate the role of SET7 in endothelial dysfunction, we generated a tamoxifen-inducible, endothelium-specific *Setd7* knockout mouse model (*Setd7*EC-KO, **Figure S1** and **Figure 2**). Thereafter, WT and *Setd7*EC-KO mice were fed STD or HFD for 12 weeks (**Figure 2A**). Consistent with its lysine methyltransferase role, the endothelium from HFD-WT mice showed higher levels of H3K4me1 and Kme1 compared with STD littermates, whereas these marks were significantly reduced in the endothelium of *Setd7*EC-KO mice regardless of diet (**Figure S4**). Both WT and *Setd7*EC-KO mice developed similar HFD-induced metabolic alterations including increased body weight, blood glucose, insulin, triglycerides, free fatty acids (FFAs), total and LDL/VLDL cholesterol levels, as well as reduced HDL cholesterol values (**Figure S5**). Despite this, wire myograph analyses of aortic rings revealed striking differences in vascular function between genotypes (**Figure 2B**). Endothelium-dependent relaxations to acetylcholine (Ach) were significantly reduced in aortas from WT-HFD mice compared with WT-STD controls (**Figure 2B**, upper panel). Interestingly, *Setd7*EC-KO mice exposed to HFD were protected from endothelial dysfunction, as their aortas exhibited significantly greater relaxations than those from WT-HFD animals (**Figure 2B**, upper panel). On the contrary, endothelium-independent relaxations to sodium nitroprusside (SNP) did not differ among groups (**Figure 2B**, lower panel).

**Figure 2.**
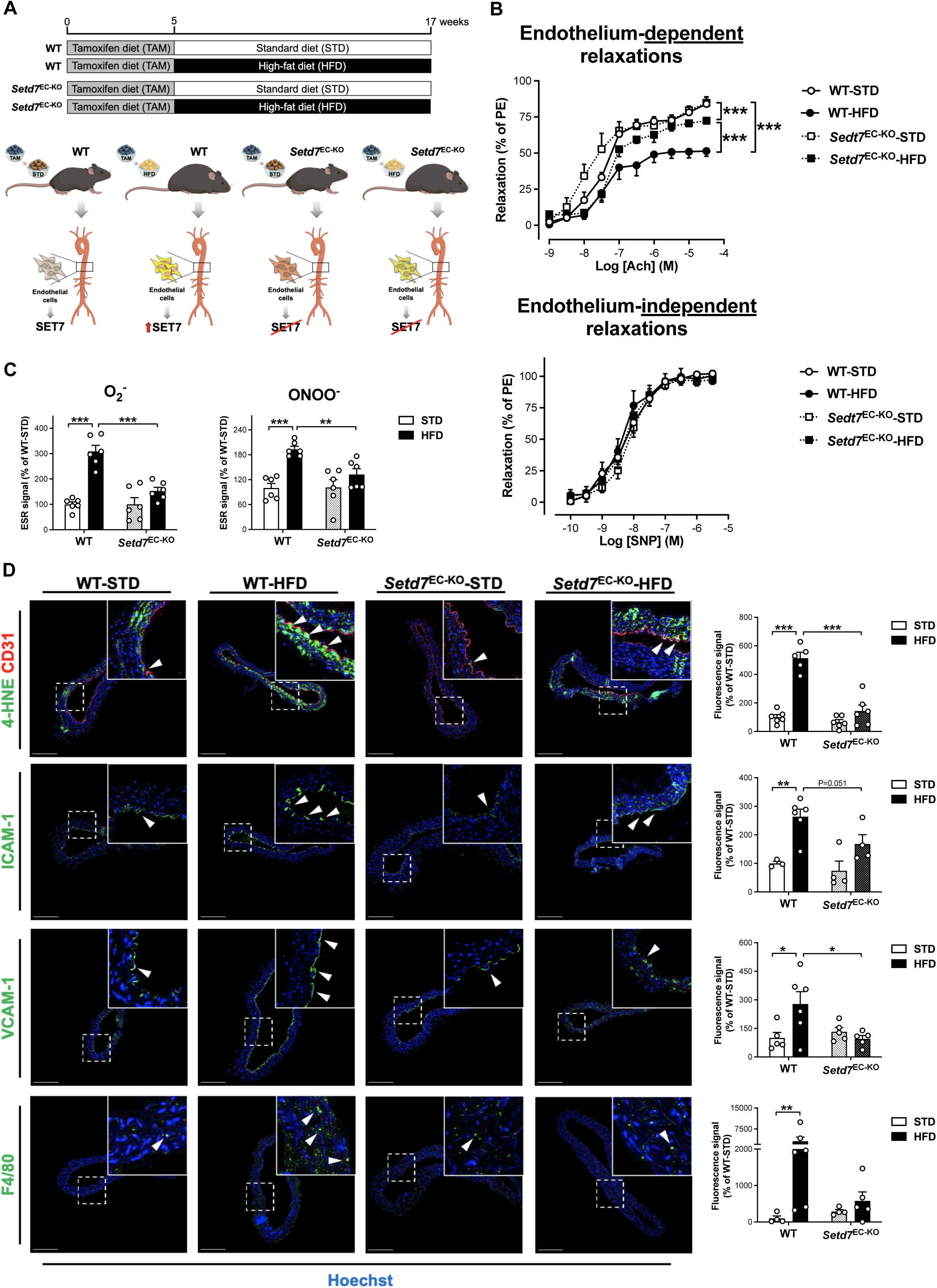
Endothelium-specific deletion of *Setd7* preserves endothelial function by attenuating oxidative stress and inflammation. **A**, Schematic representation of the experimental animal design. **B**, Endothelium-dependent and -independent relaxations, **C**, levels of reactive oxygen species (ROS), superoxide anion (O2-) and peroxinitrite (ONOO^-^), and **D**, representative confocal images and quantification of 4-HNE, ICAM-1, VCAM-1, and F4/80 (green) in aortas from WT and *Setd7*EC-KO mice fed with standard diet (STD) or high-fat diet (HFD). The endothelium was visualized by CD31 immunostaining (red) and nuclei were counterstained with Hoechst (blue). Scale bar = 200 μm. *P<0.05, **P<0.01, ***P<0.001.

To determine whether the endothelial protective effects of *Setd7* deletion are associated with reduced oxidative stress and inflammation, specific markers were analysed in aortas from the four animal groups (**Figures 2C** and **D**). Consistent with the vascular functional data (**Figure 2B**), levels of ROS, such as superoxide anion (O2-) and peroxynitrite (ONOO^-^) as well as the oxidative stress marker 4-HNE, were markedly elevated in the endothelium of WT but substantially blunted in *Setd7*EC-KO mice in response to HFD (**Figures 2C** and **D**, upper panel). Accordingly, the impaired endothelium-dependent relaxations observed in WT-HFD aortas were restored by incubation with the ROS scavengers apocynin and tempol (**Figure S6**). Moreover, aortas from *Setd7*EC-KO obese mice displayed less vascular inflammation compared to WT-HFD mice, as shown by decreased expression of the adhesion molecules ICAM-1 and VCAM-1 and lower levels of the macrophage marker F4/80 (**Figure 2D**, lower panels).

### Hyperglycemia is a Major Driver of SET7 Upregulation in Aortic Endothelial Cells and SET7 Inhibition Blunts Inflammation and Oxidative Stress

To assess how HFD-induced metabolic disturbances influence SET7 expression in aortic endothelial cells, we established in vitro models mimicking the metabolic alterations observed in vivo, namely hyperglycaemia, hyperinsulinemia, and lipid overload (**Figure S5**).

Specifically, human aortic endothelial cells (HAECs) were exposed to high concentrations of glucose (HG), insulin, fatty acids (either oleate or palmitate), and LDL. All treatmens except insulin induced increased lipid content without compromising cell viability (**Figure S7**).

Among the different stimuli tested, HG elicited the strongest upregulation of SET7 at both mRNA and protein levels, followed by oleate, palmitate, and LDL, while no changes were detected after insulin treatment (**Figures 3A** and **B**). Given these observations, we next examined the expression of a panel of relevant inflammatory and oxidative genes in HAECs exposed to each of these stimuli, either alone or in combination with SET7-specific inhibitor (SET7i) cyproheptadine^38^ (**Figure 3C** and **Figure S8**). These studies showed that SET7 inhibition markedly attenuated HG-induced expression of nearly all pro-inflammatory and pro-oxidant genes investigated (**Figure 3C**, left panels), whereas it had limited to no effect on the HG-mediated suppression of anti-inflammatory and anti-oxidant genes (**Figure 3C**, right panels). Notably, the blunting effects of SET7 inhibition on pro-inflammatory and pro-oxidant gene expressions were less pronounced under oleate, palmitate, or LDL stimulation as compared to HG (**Figure S8**), highlighting HG as a dominant driver of SET7-dependent inflammatory and oxidative responses.

**Figure 3.**
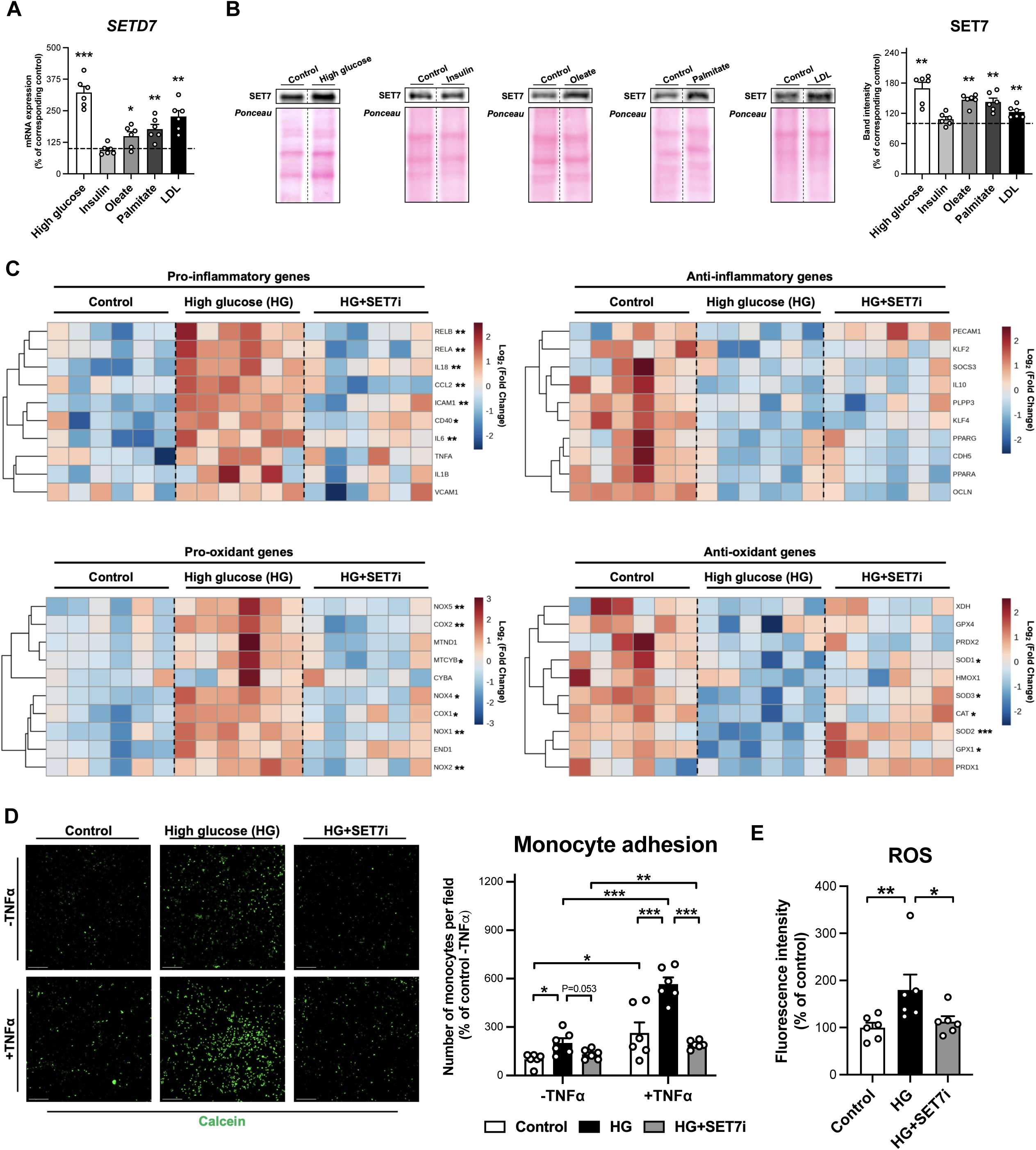
Hyperglycemia is a major driver of SET7 upregulation in aortic endothelial cells and SET7 inhibition blunts inflammation and oxidative stress. **A**, mRNA levels of *SETD7* and **B**, representative blots and quantification of SET7 in human aortic endothelial cells (HAECs) exposed to high glucose (HG, 25 mM), insulin (100 nM), oleate (250 μM), palmitate (250 μM), or LDL (20 nM) for 24 hours. **C**, Hierarchical clustering dendrogram heatmap analysis of pro-/anti-inflammatory and pro-/anti-oxidant gene expression in HAECs exposed to control, HG, or HG + SET7-specific inhibitor (SET7i) cyproheptadine (10 μM) for 24 hours. The scale in the colour bar represents -Log2 (Fold Change). The expression of genes that were significantly altered by HG and restored in the presence of SET7i are marked with asterisks corresponding to the P values indicated below. **D**, Representative confocal images and quantification of monocytes (stained with calcein, green) adherent to HAECs exposed to HG alone or with SET7i for 24 hours in the presence and in the absence of TNFα (5 nM). Scale bar = 200 μm. **E**, Reactive oxygen species (ROS) production in HAECs exposed to HG alone or with SET7i for 24 hours. *P<0.05, **P<0.01, ***P<0.001.

In line with these transcriptional findings, inhibition of SET7 significantly reduced HG- induced endothelial activation, as shown by a lower number of adherent monocytes following TNFα challenge (**Figure 3D**), and the restoration of ROS production to control levels (**Figure 3E**). Importantly, SET7 silencing (**Figures S9A** and **S9B**) recapitulated these protective effects, reversing the expression of several pro-inflammatory and pro-oxidant genes, monocyte adhesion, and ROS generation in HAECs exposed to HG (**Figures S9C-E**).

Notably, SET7 can both modulate cellular function directly, via its methyltransferase activity, and indirectly through competition with other post-translational modifying enzymes (e.g., acetyltransferases or demethylases)^48^. To distinguish between these actions, we next tested whether its catalytic activity was directly responsible for the observed pathological phenotypes (**Figure S10**). Thus, HAECs were transfected with plasmids encoding either catalytically active SET7 (SET7-WT) or an inactive mutant (SET7-MT)^4^ (**Figures S10A** and **S10B**). Interestingly, overexpression of SET7-WT, but not SET7-MT, strongly enhanced the expression of multiple pro-inflammatory and pro-oxidant genes, increased monocyte adhesion, and ROS production (**Figures S10C-E**).

### SET7-mediated H3K4me1 Drives Endothelial Inflammation and Oxidative Stress Through mRNA Splicing Gene Activation

To identify H3K4me1-dependent targets of SET7 in endothelial cells under hyperglycemic conditions, we performed proteomic profiling of HAECs exposed to control, HG, or HG+SET7i (**Figure 4A**). As shown in **Figure 4B**, HG treatment significantly induced the expression of 1,171 proteins (left panel), whereas the abundance of 920 proteins was reduced in the presence of SET7i (right panel). Overlap of these datasets revealed 668 proteins whose expression was induced in HG and reversed by SET7i, thereby representing putative SET7- regulated histone methylation targets (**Figure 4C**). Pathway analysis of these proteins using Reactome^49^ showed that the four most significantly overrepresented categories are related to RNA metabolism, including 33 proteins involved in mRNA splicing (**Figure 4D** and **Table S2**). Among them, proteins that participate in each step of the splicing process^50^ were found: assembly of the splicing machinery (i.e., spliceosome; 10 proteins), its activation (8 proteins), splicing catalysis (12 proteins), and spliceosome disassembly (3 proteins) (**Figure 4E**).

**Figure 4.**
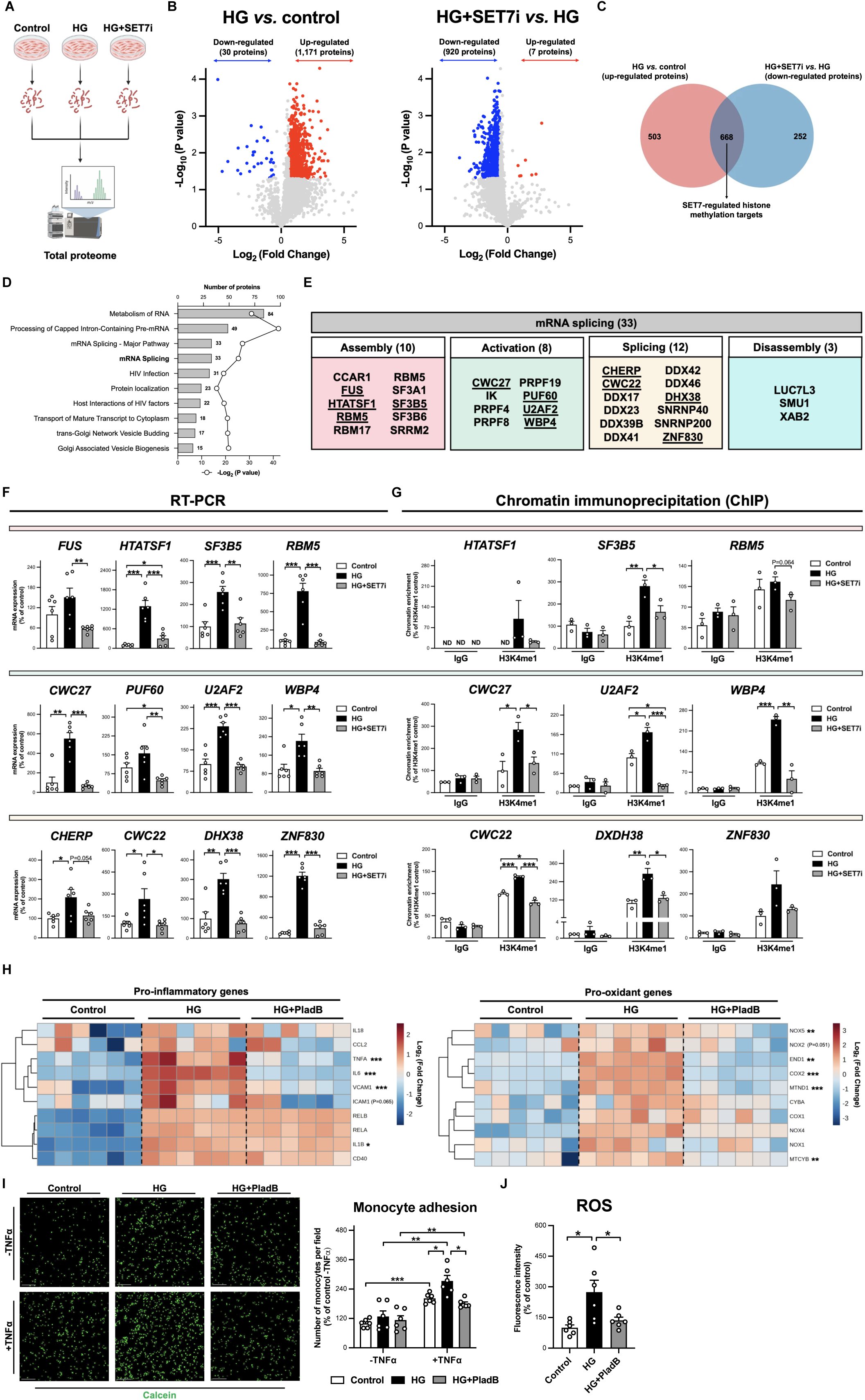
SET7-mediated H3K4me1 drives endothelial inflammation and oxidative stress through mRNA splicing gene activation. **A**, Schematic representation of the experimental design. **B**, Volcano plots representing the total proteome of HAECs comparing HG *vs.* control and HG+SET7i *vs.* HG. The x-axis shows the Log2 (Fold Change) and the y-axis the corresponding -Log10 (P value) of each identified protein. Significantly down- and upregulated proteins [Log2 (Fold Change) > ±1.5 and P value < 0.05] are in blue and red, respectively, whereas non significant proteins are in grey. **C**, Venn diagramn showing overlap of proteins that are significantly upregulated in HAECs by HG and downregulated in response to SET7i. **D**, Top ten over-represented pathways related to proteins identified as putative SET7-regulated histone methylation targets according to Reactome^49^. Numbers indicate the number of identified proteins in the pathway, whereas circles indicate −Log2 (P value). **E**, Proteins identified in the “mRNA splicing” pathway, grouped according to their primary role in each step of the process: spliceosome assembly (pink), activation (green), splicing (yellow), or disassembly (blue). Underlined proteins were selected for further analyses due to their critical participation in each step. **F**, mRNA levels of selected genes in each step: spliceosome assembly (*FUS*, *HTATSF1*, *RBM5*, *SF3B5*; pink), activation (*CWC27*, *PUF60*, *U2AF2*, *WBP4*; green), and splicing (*CHERP*, *CWC22*, *DHX38*, ZNF830; yellow) in HAECs exposed to control, HG, or HG+SET7i for 24 hours. **G**, H3K4me1 enrichment in the promoters of selected genes in each step: assembly (*HTATSF1*, *RBM5*, *SF3B5*; pink), activation (*CWC27*, *U2AF2*, *WBP4*; green), and splicing (*CWC22*, *DHX38*; yellow) in HAECs exposed to control, HG, or HG+SET7i for 24 hours. **H**, Hierarchical clustering dendrogram heatmap analysis of pro- inflammatory and pro-oxidant gene expression in HAECs exposed to control, HG, or HG+SET7i for 24 hours. The scale in the colour bar represents -Log2 (Fold Change). Genes whose expression is significantly altered in HG and significantly or tend to be restored by SET7i are indicated with asterisks or the corresponding P value, respectively. **I**, Representative confocal images and quantification of monocytes (stained with calcein, green) adherent to HAECs exposed to control, HG, or HG + splicing inhibitor Pladienolide-B (PladB) for 24 hours in the presence and in the absence of TNFα (5 nM). Scale bar = 200 μm. **J**, Reactive oxygen species (ROS) production in HAECs exposed to control, HG, or HG+PladB for 24 hours. *P<0.05, **P<0.01, ***P<0.001.

To validate these findings at the transcript level, we examined the expression of the most relevant splicing-related genes, as defined by the spliceosome atlas^50^. In agreement with our proteomic data, HG significantly upregulated the mRNA content of genes involved in spliceosome assembly (*HTATSF1*, *RBM5*, *SF3B5*), activation (*CWC27*, *U2AF2*, *WBP4*), and catalysis (*CWC22*, *DHX38*, *ZNF830*), all of which were significantly normalised by SET7 inhibition (**Figure 4F**). Similar trends were observed for *FUS* (involved in spliceosome assembly), *PUF60* (activation), and *CHERP* (splicing), although these changes did not reach statistical significance (**Figure 4F**). To determine whether the transcription of these genes is directly regulated via SET7-mediated H3K4me1, ChIP-qPCR assays were performed (**Figure 4G**). HAECs exposed to HG displayed significantly increased (or higher-trending) H3K4me1 enrichment at the promoter regions of all analysed genes, which was reversed by SET7 inhibition (**Figure 4G**), confirming a crucial regulatory role of SET7-mediated H3K4me1 in controlling mRNA splicing gene transcription (**Figure 4G**).

Given the established role of aberrant mRNA splicing in vascular injury and atherosclerosis^51,52^, we next investigated whether spliceosome hyperactivation induced by HG contributes to endothelial dysfunction. In fact, pharmacological inhibition of splicing using Pladienolide-B (PladB)^53,54^ attenuated HG-induced inflammatory and oxidant gene expression (**Figure 4H**), along with attenuation of both monocyte adhesion and ROS generation (**Figures 4I** and **4J**). Taken together, these results indicate that SET7-dependent H3K4me1 promotes endothelial inflammation and oxidative stress by enhancing the transcription of key components of the mRNA splicing machinery.

### SET7-Induced Methylation of eNOS at K494 Disrupts Calmodulin Binding and Reduces NO Production in Aortic Endothelial Cells

To elucidate protein targets of SET7-mediated Kme1 in HAECs, we performed Kme1 proteomics in control cells and cells treated with HG or HG+SET7i (**Figure 5A**). Kme1- modified proteins were enriched via immunoprecipitation using an in-house manufactured antibody (**Figure S3**). Importantly, methylation levels were normalised to total protein content, ensuring that observed changes reflect true alterations in methylation rather than in protein abundance. This analysis identified 18 proteins that were hyper-methylated in response to HG (**Figure 5B**, left panel) and 1,199 proteins that were hypo-methylated upon SET7 inhibition (**Figure 5B**, right panel), underscoring a broad regulatory role of SET7 in protein methylation. Intersection of these two datasets revealed 15 proteins as putative SET7 non-histone substrates (**Figure 5C**). STRING network analysis^55^ indicated that these proteins participate in diverse pathways, including NO signalling, gene transcription, cytoskeleton organisation, inflammation, vesicle transport, and mitochondrial metabolism (**Figure 5D**). Among them, NOS3 (or endothelial nitric oxide synthase, eNOS) emerged as one of the most strongly affected proteins by SET7-mediated Kme1 in response to HG (**Figures 5B** and **5D**).

**Figure 5.**
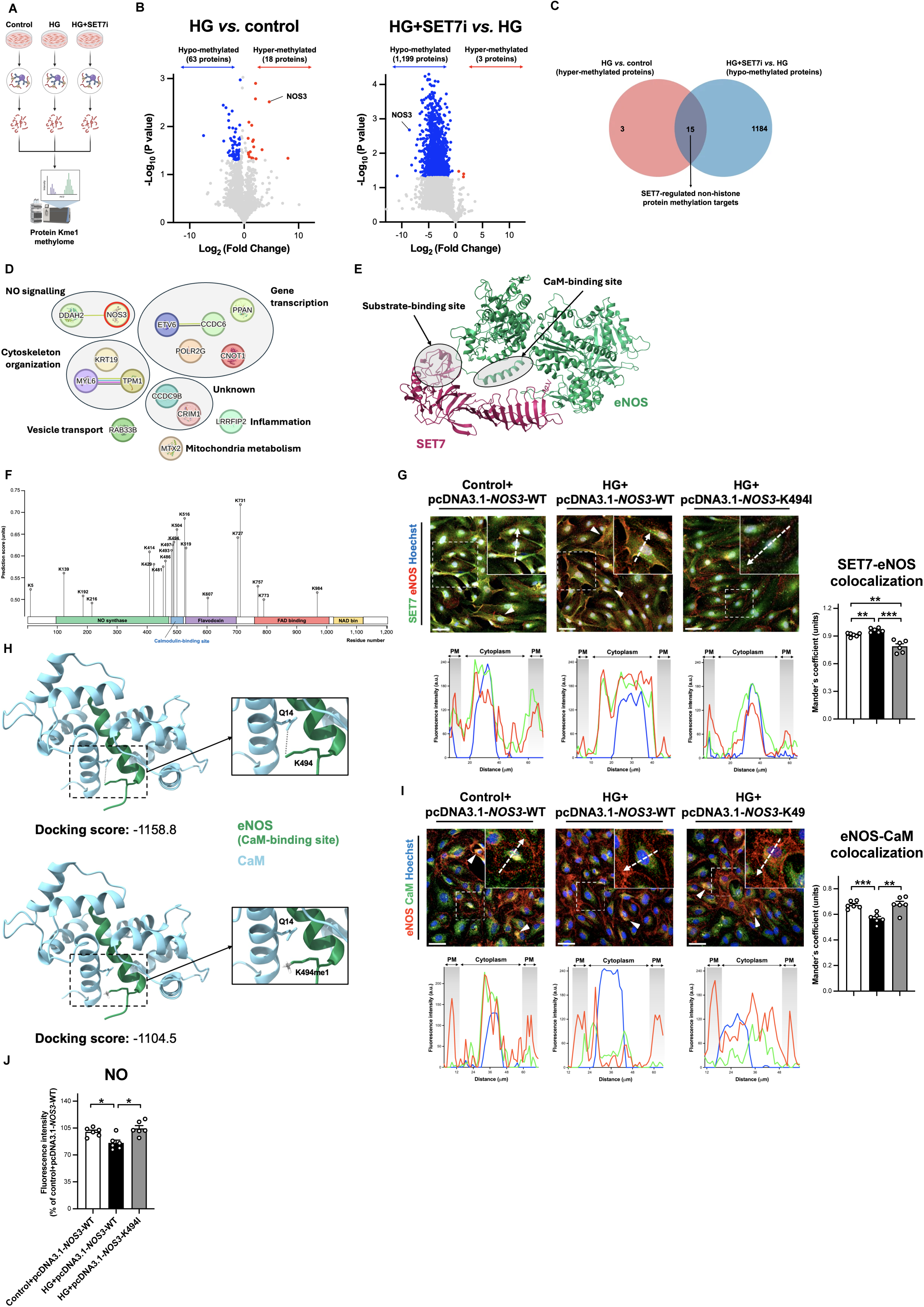
SET7-induced methylation of eNOS at K494 disrupts calmodulin binding and reduces NO production in aortic endothelial cells. **A**, Schematic representation of the experimental design. **B**, Volcano plots representing the protein Kme1 methylome of HAECs comparing HG *vs.* control and HG+SET7i *vs.* HG. The x-axis shows the Log2 (Fold Change) and the y-axis the corresponding -Log10 (P value) of each identified protein. Significantly [Log2 (Fold Change) > 1.5 and P value < 0.05] hyper- and hypo-methylated proteins are in blue and red, respectively, whereas non significant proteins are in grey. **C**, Venn diagramn showing the overlap of proteins that are significantly hyper-methylated in HAECs by HG and significantly hypo-methylated in response to SET7i. **D**, Interaction of SET7-regulated non-histone protein methylation targets according to STRING^55^. **E**, Protein-protein docking analyses of SET7 (PDB ID: 1H3I, pink) and eNOS (AlphaFold ID: AF-P29474-F1-v6, green). **F**, eNOS protein sequence and prediction scores of lysine methylation according to MethylSight^57^. **G**, Representative confocal images, quantification, and distribution plots of SET7 (green) and eNOS (red) colocalization in HAECs exposed to control or HG conditions overexpressing pcDNA3.1-*NOS3*-WT or *NOS3*-K494I. **H**, Protein-protein docking analyses and scores of the calmodulin (CaM)-binding site of eNOS (PDB ID: 1NIW, green) and CaM (blue) in normal conditions (left panel) and when K494 is mono-methylated (right panel). **I**, Representative confocal images, quantification, and distribution plots of eNOS (red) and CaM (green) colocalization in HAECs exposed to control or HG conditions overexpressing pcDNA3.1-*NOS3*-WT or *NOS3*-K494I. **J**, Nitric oxide (NO) production levels in HAECs exposed to control or HG conditions overexpressing pcDNA3.1-*NOS3*-WT or *NOS3*-K494I. P<0.05, **P<0.01, ***P<0.001. PM, plasmatic membrane; Scale bar = 50 μm.

Protein-protein docking analysis showed that the SET7 substrate-binding pocket likely aligns adjacent to the calmodulin (CaM)-binding region of eNOS (**Figure 5E**). Given the flexible hinge-like action of methyltransferases, which fold upon themselves to reach target residues^56^, this positioning suggests that SET7 methylates lysines in this region. In support of this, prediction of methylation sites using MethylSight^57^ identified a cluster of lysine residues with high methylation probability in the CaM-binding domain of eNOS, with K494 ranking the highest (**Figure 5F**). To experimentally validate this finding, we carried out site-directed mutagenesis of *NOS3* (**Figure S11A)**. Thus, we generated a *NOS3* mutant construct (i.e., *NOS3*-K494I) in which the lysine 494 codon (AAG) was replaced with isoleucine (ATT) (**Figure S11B**). Isoleucine was selected because it can not be recognised by SET7, yet it is structurally similar to lysine^58^, preserving the overall tridimensional conformation of eNOS (**Figure S11C**). Next, HAECs were transfected with WT- or K494I-eNOS plasmids, resulting in comparable overexpression magnitudes (**Figure S11D**). Confocal microscopy studies revealed that SET7 and eNOS colocalization increased under HG in cells overexpressing WT- eNOS, but was markedly diminished in those cells expressing NOS3-K494I (**Figure 5G**), confirming SET7-driven methylation of eNOS at this specific residue within the CaM-binding site.

To investigate the relevance of eNOS methylation at K494, structural modelling of eNOS, with or without K494 methylation, interacting with CaM was generated (**Figure 5H**). The CaM-methylated eNOS complex exhibited a less negative docking score than with the unmethylated eNOS, indicating lower binding affinity (**Figure 5H**, lower panel). Such a decrease could be related to the loss of stabilising hydrogen bonds between CaM glutamine 14 (Q14) and K494-eNOS (**Figure 5H**, zoom boxes). Accordingly, HG-induced reduction in eNOS-CaM colocalization was normalised in HAECs overexpressing the eNOS-K494I variant (**Figure 5I**). Functionally, since CaM binding is essential for eNOS activation and NO synthesis^59^, we measured NO levels in HAECs transfected with the two forms of eNOS, and found that in cells expressing *NOS3*-K494I under HG conditions NO levels were restored, whereas eNOS-WT-expressing cells continued to display impaired NO generation (**Figure 5J**). These results explicitly associate SET7-mediated methylation of eNOS at K494 to HG-induced impairment of NO production in HAECs.

### SET7-Dependent Activation of mRNA Splicing and Impairment of eNOS-CaM Interaction in Arteries from Patients with Hyperglycemia

Finally, to explore whether the molecular alterations identified in the WT-HFD mouse model and HG-exposed HAECs extend to human vascular disease, we analysed internal mammary arteries (IMAs) from patients undergoing coronary artery bypass graft surgery (**Figure 6**). Patients were stratified into two groups, according to their glycaemic status, based on the American Diabetes Association criteria60. Subjects with normoglycemia (NG; HbA1c < 39 mmol/mol), and with impaired fasting glucose or T2D (IFG/T2D; HbA1c ≥ 39 mmol/mol). No significant differences were found between the two groups in any other clinical or biochemical variables, except for the higher fasting glucose, HbA1c, and lower HDL cholesterol observed in IFG/T2D individuals (**Table 1**).

**Figure 6.**
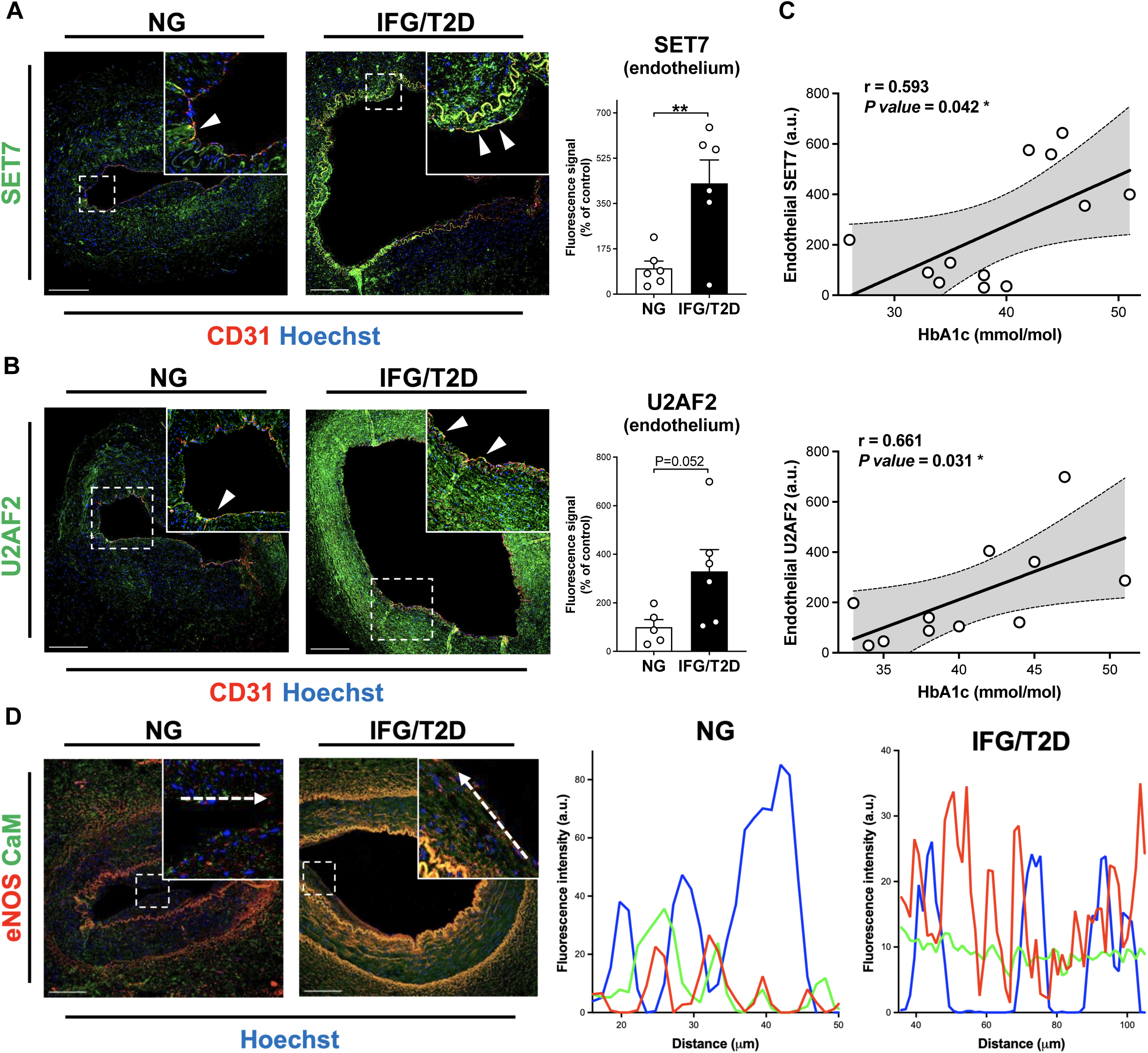
SET7-dependent activation of mRNA splicing and impairment eNOS-CaM interaction in arteries from patients with hyperglycemia. Representative confocal images and quantification of **A**, SET7 and **B**, U2AF2 (green) in the endothelium layer of internal mammary arteries from patients with normoglycemia (NG) or impaired fasting glucose/type 2 diabetes (IFG/T2D). The endothelium was visualized by CD31 immunostaining (red) and nuclei were counterstained with Hoechst (blue). **C**, Linear regression analysis between endothelial SET7 (upper panel) or U2AF2 (lower panel) expression levels (arbitrary units, a.u.) and circulating levels of HbA1c. *P<0.05. **D**, Representative confocal images and distribution plots of eNOS (red) and CaM (green) colocalization in internal mammary arteries from patients with normoglycemia (NG) or impaired fasting glucose/type 2 diabetes (IFG/T2D). Nuclei were counterstained with Hoechst (blue). Scale bar = 200 μm.

In accordance with the results observed in HFD-fed mice (**Figure 1C**) and the HG model in HAECs (**Figure 3B**), confocal microscopy analyses revealed a marked increase in endothelial SET7 expression in arteries from IFG/T2D vs. NG patients. In parallel, endothelial levels of U2AF2, a spliceosome-associated factor that we found to be strongly regulated by both HG exposure and SET7 inhibition in HAECs (**Figures 4F** and **4G**), were also clearly elevated in IFG/T2D arteries, showing a strong trend towards statistical significance in comparison to NG specimens (**Figure 6B)**. Importantly, endothelial SET7 and U2AF2 expression levels showed a significant positive correlation with circulating HbA1c values (**Figure 6C**), supporting a close association between hyperglycemia and activation of the SET7-splicing programme in human vasculature. In addition, endothelial SET7 expression was also positively correlated with body mass index and fasting glucose, and both SET7 and U2AF2 expressions with circulating triglyceride concentrations (**Table S3**). Moreover, given our previous findings linking SET7 to impaired eNOS-CaM interaction under hyperglycemic conditions (**Figure 5I**), we next examined the spatial distribution of these proteins in human arteries. In vessels from NG patients, CaM immunosignal extensively colocalised with eNOS within the endothelial layer. By contrast, this pattern was disrupted in arteries from IFG/T2D individuals (**Figure 6D**). Collectively, these results indicate that the molecular signatures identified in experimental models, including HG-associated SET7 upregulation (**Figure 1**), activation of an mRNA splicing program (**Figure 4**), and disruption of the eNOS-CaM interaction (**Figure 5**), are recapitulated in arteries from patients with vascular disease and hyperglycemia. These observations support the translational relevance of SET7-dependent endothelial dysfunction and identify this lysine methyltransferase as a key mediator linking elevated plasma glucose levels to vascular injury in human cardiometabolic disease.

## DISCUSSION

Herein, we provide experimental evidence demonstrating selective upregulation of endothelial SET7 in response to diet-induced metabolic derangements, particularly hyperglycemia. Both endothelial-specific deletion and pharmacological inhibition of SET7 preserve endothelium-dependent vasoreactivity and attenuate vascular oxidative stress and inflammation. Mechanistically, we uncover two novel complementary pathways through which SET7 promotes endothelial dysfunction in this setting: (i) H3K4me1-dependent transcriptional activation of spliceosome components promoting pro-inflammatory and pro-oxidative phenotypes, and (ii) direct methylation of eNOS at K494 that impairs CaM binding and reduces NO bioavailability. Interestingly, these mechanisms are also activated in arteries from ASCVD patients with hyperglycemia, supporting their clinical relevance. Taken together, these findings position SET7 as an important epigenetic and post-translational mediator linking metabolic imbalance to vascular injury.

We have previously shown that SET7 expression is increased in circulating monocytes from individuals with T2D, contributing to vascular dysfunction^61^. The present study substantially extends these observations, showing that, within the vascular wall, SET7 expression is specifically induced in the endothelium and that this event is causally required for the occurrence of diet-induced endothelial dysfunction in vivo. In particular, using a unique endothelial-specific *Setd7* knockout mouse model, we found that the loss of SET7 preserves endothelium-dependent relaxations in aortas from HFD-fed mice, as assessed by wire myography. Importantly, endothelial *Setd7* deletion did not affect the metabolic impairment induced by HFD, placing SET7 as a cell-autonomous regulator and biomarker of endothelium damage.

This work also confirms and expands previous reports from our and other groups implicating SET7 in vascular oxidative stress and inflammation^28^,^61–63^. Specifically, the quantification of O2- and ONOO^-^ by ESR, together with the complementary assessment of the oxidative stress marker 4-HNE in mouse aortas, support the conclusion that endothelial SET7 deletion markedly attenuates vascular oxidative stress in vivo. However, we found that although antioxidant treatment with apocynin or tempol significantly improved endothelial function in HFD aortas, it did not fully restore acetylcholine-induced relaxations. This partial rescue suggests the involvement of additional ROS-independent mechanisms. In line with this observation, the reduced expression of adhesion molecules and decreased macrophage infiltration observed in aortas from obese *Setd7*EC-KO mice substantiates a driving role of SET7 in endothelial inflammation. These findings are particularly noteworthy given that excessive ROS production and monocyte adhesion are hallmark features of endothelial dysfunction and are widely regarded as central early events in atherogenesis^64,65^. Although atherosclerosis was not directly examined here, SET7 has recently been shown to be upregulated in human carotid atherosclerotic lesions and in the aorta of ApoE-/- mice with established atherosclerosis^28^. Moreover, long-term systemic blockade of SET7 catalytic function reduces atherosclerotic plaque formation in these mice^28^, suggesting that the endothelial protection exerted by SET7 deletion and inhibition identified in the present study is likely to have important implications for the progression of ASCVD.

Another strength of our study is the dissection of upstream metabolic cues that drive SET7 upregulation in endothelial cells. Although enhanced SET7 expression has been documented in individuals with diabetes^61,66^, the mechanisms governing its induction in the endothelium under pathological conditions remain poorly defined. To address this gap, we employed in vitro models that recapitulate key metabolic features of obesity and insulin resistance observed in our HFD mouse model and in individuals with obesity and diabetes. These included hyperglycemia, hyperinsulinemia, hyperlipidemia, modelled using either saturated (palmitate) or unsaturated (oleate) fatty acids, and hypercholesterolemia (LDL cholesterol). Notably, high glucose emerged as the dominant stimulus driving SET7 upregulation in HAECs, with fatty acids and LDL cholesterol producing more modest but still significant effects. While hyperglycemia-induced SET7 expression has been reported before by us and others^61,63,67^, our current findings expand these observations by implicating SET7 in broader metabolic stress responses. This aligns with the notion that intensive glycemic control alone is often insufficient to fully reverse endothelial dysfunction in diabetes, as vascular damage arises from a complex, multifactorial environment that goes beyond hyperglycemia^68^. Intriguingly, SET7 induction occurred concomitantly with the accumulation of intracellular lipid droplets, suggesting a previously unrecognised link between SET7 expression and endothelial lipid handling. Consistent with this, SET7 has been shown to increase fatty acid synthase (FASN) expression, thereby enhancing FA metabolism in cancer cells^69^. However, pharmacological inhibition of SET7 failed to fully restore lipid-mediated molecular changes in HAECs, suggesting that additional molecular determinants contribute to these effects. Future studies will establish whether other pathogenic processes causing endothelial dysfunction in metabolic disease (e.g., fibrosis, hypoxia, altered shear stress) may also modulate SET7 expression in these cells.

In line with our in vivo genetic deletion studies, pharmacological inhibition of SET7 robustly reversed hyperglycemia-induced changes in HAECs. Specifically, we employed cyproheptadine, a clinically approved drug that binds to the substrate-binding pocket of SET7 and inhibits its enzymatic activity by competing with the methyl group acceptor^70^. Cyproheptadine exclusively impedes SET7-dependent methylation without affecting other SET-domain methyltransferases^70^, and has been successfully used to interrogate SET7 function in other diseases^71–73^, supporting its suitability as a pharmacological tool. Importantly, SET7 inhibition in this study not only restored HG-induced expression of pro-inflammatory and pro-oxidant genes but also led to biologically relevant functional improvements, including diminished monocyte adhesion and reduced endothelial ROS generation. These effects were consistently observed across multiple complementary experimental approaches, including siRNA-mediated silencing and overexpression studies. Gain-of-function experiments were specifically employed to establish direct causality and delineate the contribution of SET7 enzymatic activity. This distinction is particularly important because SET7 has been reported to exert catalysis-independent functions as well, an aspect that has been systematically overlooked in prior studies examining SET7’s role in disease. Indeed, the N-terminal MORN repeat domains of SET7 facilitate substrate docking and mediate non-enzymatic interactions with transcription factors, chromatin-associated proteins, and transcriptional co-regulators such as p300/CBP, SWI/SNF, and SRF, all of which are known to influence gene regulation^74–76^. Notably, we show that overexpression of catalytically active SET7, but not its inactive mutant, markedly exacerbates inflammation and oxidative stress in HAECs, thereby excluding a dominant role for catalysis-independent mechanisms and directly implicating SET7 methyltransferase activity as the principal driver of the observed alterations. Together, these results highlight the translational relevance of targeting SET7 and indicate that pharmacological inhibition of its enzymatic activity may represent a viable therapeutic strategy to mitigate endothelial dysfunction in cardiometabolic disease, as it has been proposed for other pathological settings^77^.

SET7 is a well-established histone lysine methyltransferase that catalyses H3K4me1, a chromatin mark associated with transcriptionally active regions and permissive gene expression programs^78^. A major conceptual advancement of our study is the demonstration that SET7-driven H3K4me1 enhances the transcription of spliceosome-related genes, thereby directly linking epigenetic reprogramming to dysregulation of mRNA splicing in endothelial cells. Proteomic analyses revealed that SET7 activity accounted for the HG-induced expression of a substantial proportion of proteins (>600 proteins), including key representative components spanning all major steps of the splicing process. Importantly, these changes were mirrored at the mRNA level, proving its transcriptional control. SET7-dependent enrichment of H3K4me1 at the promoters of these splicing-related genes was further confirmed by ChIP experiments, establishing a direct mechanistic link between SET7 activity and spliceosome gene transcription. Deleterious mRNA splicing has recently emerged as an important pathogenic mechanism in atherosclerosis and vascular inflammation^79,80^, yet the upstream signals governing activation of the splicing machinery have not been well defined. Our findings place SET7 at the centre of this process, providing a mechanistic explanation for how metabolic stress reshapes the endothelial transcriptome and proteome. Specifically, HG-induced SET7 activation promotes overexpression of splicing factors, which in turn contributes to inflammatory and oxidative endothelial phenotypes. Consistent with this model, pharmacological inhibition of mRNA splicing using the spliceosome inhibitor PladB effectively reversed inflammatory and oxidative stress responses, supporting a causal role for spliceosome hyperactivation in endothelial dysfunction. mRNA splicing is a fundamental mechanism that enables cells to expand proteomic diversity and dynamically adapt gene function in response to environmental cues. Increasing evidence indicates that mRNA splicing plays a critical role in cardiovascular biology by modulating endothelial signalling, inflammatory responses, and vascular remodelling through the generation of disease-specific isoforms^79,80^. Under pathological conditions, excessive or promiscuous spliceosome activity can lead to aberrant exon inclusion or exclusion, cryptic splice-site usage, and the production of truncated protein isoforms that exacerbate cellular dysfunction. In this regard, our data support a model in which hyperactivation of the splicing machinery under HG conditions may contribute, at least in part, to pathogenic isoform switching that drives endothelial dysfunction in metabolic disease. Together, these findings add to the current repertoire of SET7-mediated H3K4me1 targets beyond classical inflammatory gene networks and establish splicing regulation as a novel epigenetic output of SET7 activity in the endothelium.

In addition to histone modifications, SET7 mono-methylates a broad range of non-histone proteins (Kme1)^10,11^, including p53^12,13^, NF-kB p65^14,15^, STAT3^16^, YAP^17,18^, DNMT1^19,20^, FOXO3^21,22^, SIRT1^23^, and SUV39H1^24^. Despite these individual reports, a systematic evaluation of SET7 non-histone substrates had not been performed so far. This limitation largely stems from the technical challenges associated with detecting mono-methylation, a chemically subtle modification that is difficult to enrich and discriminate by mass spectrometry^81^. Moreover, the lack of highly specific and validated Kme1 antibodies has historically precluded comprehensive profiling approaches. In the present study, we overcame these limitations by combining an in-house-generated Kme1-specific antibody with unbiased proteomic analysis to generate, to our knowledge, the first lysine mono-methylome of human endothelial cells. Mapping the endothelial Kme1 landscape is particularly important given the central role of endothelial cells in vascular homeostasis and their sensitivity to metabolic and inflammatory stressors that drive ASCVD. Our data provide a previously unexplored regulatory layer of endothelial protein function, unmasking Kme1 as a widespread modification in this cell type. Interestingly, SET7 inhibition led to a marked reduction in the methylation of more than 1,000 proteins in HAECs, highlighting a far more extensive role for SET7 in defining the human lysine mono-methylome than recognized before. These findings position SET7 as a central regulator of non-histone protein methylation and suggest that its activity broadly influences endothelial signaling networks rather than acting on a limited set of substrates.

Among the SET7-dependent targets identified under HG conditions, we uncovered a novel non-histone substrate with direct implications for endothelial function: eNOS methylation at K494. eNOS is the primary enzymatic source of NO in endothelial cells, and it is essential for the regulation of vascular tone, platelet aggregation, and inflammation. To the best of our knowledge, this is the first report describing methylation of eNOS, adding to the extensive repertoire of post-translational modifications, mostly phosphorylation, that have been shown to regulate eNOS activity for decades82. Using protein-protein docking analyses and site-directed mutagenesis, we demonstrate that methylation of eNOS at K494, located within the CaM-binding domain, disrupts CaM-eNOS interaction, a critical step for eNOS activation^82^, thereby reducing NO production. Given that diminished NO bioavailability is a hallmark of endothelial dysfunction^83^, our findings establish a direct mechanistic link between SET7 activity and the abnormal vascular phenotype observed in high cardiometabolic risk. Previous studies have shown that K494 and its neighbouring amino acids, such as T495, are essential for eNOS function^84,85^. Notably, the CaM-binding region and adjacent domains of eNOS contain multiple lysine residues with a high predicted probability of methylation. It is therefore plausible that additional lysine methylation events occur simultaneously, further exacerbating the impairment of CaM binding and NO synthesis. This hypothesis warrants further investigation. Moreover, our proteomic analysis identified a broad spectrum of SET7- dependent protein methylation events, of which only NOS3 was functionally validated here. Future work should explore the contribution of Kme1 of other candidate proteins to vascular injury.

The clinical relevance of our results was reinforced by the study of internal mammary arteries (IMAs) from patients with NG or IFG/T2D undergoing coronary artery bypass grafting. Consistent with our experimental models, endothelial SET7 and U2AF2 expression were associated with chronic hyperglycaemia and uncoupled eNOS–CaM was present in arteries from IFG/T2D subjects. These findings suggest that activation of SET7-dependent pathways may also occur in the human diabetic vasculature. While IMAs serve as a clinically valuable and widely used source of human vascular tissue, their relative resistance to atherosclerosis compared to other arterial beds^86^ represents a potential limitation in capturing plaque development. Nonetheless, the pathological features examined in this work, namely endothelial dysfunction, oxidative stress, and inflammation, have been consistently documented in IMAs from patients with diabetes^87,88^, supporting the translational value and validity of this model.

In conclusion, this study identifies SET7 as a central mediator connecting metabolic impairment to vascular dysfunction, through at least two convergent mechanisms: H3K4me1- mediated transcriptional control of mRNA splicing genes and non-histone mono-methylation of key endothelial proteins such as eNOS. Our work provides compelling evidence for SET7 inhibition as a promising strategy to safeguard endothelial integrity under elevated cardiometabolic risk. Although the widespread transcriptional and post-translational influence of this lysine methyltransferase warrants careful target considerations, such intervention could offer a complementary approach to current treatments by directly addressing the vascular consequences of metabolic disease.

## Affiliations

Division of Cardiology, Department of Medicine-Solna, Karolinska Institutet; Stockholm, Sweden (J.S.-C., G.F., J.Z., O.K., F.C.). Biofilms Research Center for Biointerfaces, Department of Biomedical Science, Faculty of Health and Society, Malmö University, Sweden. Current affiliation: Acrivon AB; Lund, Sweden (M.E.J.). Department of Medical Biochemistry and Biophysics, Karolinska Institutet; Stockholm, Sweden. Current affiliation: Department of Medical Biochemistry and Biophysics, Karolinska Institutet; Stockholm, Sweden (A.V.). Division of Cardiovascular Medicine, Department of Medicine-Solna, Karolinska Institutet and the Center for Molecular Medicine, Karolinska University Hospital, Stockholm, Sweden (C.L., C.E.H.). Division of Cardiothoracic Surgery, Department of Molecular Medicine and Surgery, Karolinska Institutet, Stockholm, Sweden (E.C.). Heart, Vascular and Neuro Theme, Cardiology Unit, Karolinska University Hospital; Stockholm, Sweden (F.C., O.K.).

## Acknowledgements

The authors gratefully acknowledge Magdalena Paolino for sharing THP-1 cells as well as Pauline Döttling and David Sánchez-Fernández from Department of Medicine-Solna (MedS), Karolinska Institutet for their technical help.

## Author Contributions

Julia Sánchez-Ceinos: Conceptualization, methodology, software, validation, formal analysis, investigation, resources, data curation, writing – original draft, writing – review and editing, visualization, supervision, project administration, funding acquisition. Georgios Filis: Methodology, formal analysis, investigation, writing – review and editing. Jingyi Zhang: Methodology, formal analysis, investigation, writing – review and editing. Magnus E. Jakobsson: Resources (Kme1 antibody), writing – review and editing, visualization. Ákos Végvári: Methodology (proteomics), formal analysis, investigation, data curation, writing – review and editing, visualization. Cheukyau Luk: Methodology (internal mammary artery samples), writing – review and editing. Emelie Carlestål: Resources (internal mammary artery samples), writing – review and editing. Carolina E. Hagberg: Resources (internal mammary artery samples), writing – review and editing. Oskar Kövamees: Resources (internal mammary artery samples), writing – review and editing. Francesco Cosentino: Supervision, resources, writing – review and editing, project administration, funding acquisition.

## Supplemental Materials

Supplemental Methods

Supplemental Figures (Figures S1-S11)

Supplemental Tables (Tables S1-S4)

Major Resources Table

ARRIVE Guidelines

Uncropped Western Blots and Original Images

References 1-24

## Nonstandard Abbreviations and Acronyms

ASCVD: Atherosclerotic cardiovascular disease
T2D: Type 2 diabetes
H3K4me1: Mono-methylation of histone H3 at lysine 4
HFD: High-fat diet
IMAs: Internal mammary arteries
Kme1: Mono-methylation of lysine
STD: Standard diet
HAECs: Human aortic endothelial cells
HG: High glucose
PladB: Pladienolide-B
SET7i: SET7 inhibitor

## Novelty and Significance What is Known?

- Lysine methyltransferase SET7 regulates gene expression through H3K4me1 and modifies the function of selected non-histone proteins through Kme1.
- Endothelial dysfunction is a hallmark of cardiometabolic disease, but the specific role of SET7 and its downstream molecular targets in the endothelium remain poorly defined.
- Aberrant mRNA splicing and reduced NO contribute to endothelial inflammation and oxidative stress, but their upstream molecular drivers under metabolic stress are not elucidated.

## What New Information Does This Article Contribute?

- Endothelial-specific deletion or pharmacological inhibition of SET7 protects against high-fat diet– and hyperglycemia-induced endothelial dysfunction by reducing oxidative stress and inflammation.
- SET7 promotes endothelial dysfunction by combining epigenetic activation of spliceosome genes that drive inflammation and oxidative stress with post-translational inhibition of eNOS activity through impaired calmodulin binding.
- Molecular signatures of SET7 induction, increased spliceosome genes, and disruption of the eNOS–calmodulin interaction are present in arteries from patients with ASCVD and hyperglycemia, supporting the translational relevance of these mechanisms.

Cardiometabolic disorders, including obesity and type 2 diabetes, are major drivers of atherosclerotic cardiovascular disease, largely through the development of endothelial dysfunction for which specific mechanism-based therapies are unavailable. Here, we identify lysine methyltransferase SET7 as a novel regulator linking metabolic stress to vascular injury. Using endothelial-specific *Setd7* knockout mice exposed to high-fat diet and human aortic endothelial cells challenged with high glucose, we show that genetic deletion and pharmacological inhibition of SET7 preserve endothelial function regardless of persistent systemic metabolic abnormalities. These findings indicate that SET7 acts downstream of metabolic cues in the endothelium. Then, by proteomic and chromatin analyses, we demonstrate that SET7 causes endothelial dysfunction through both epigenetic and post-translational mechanisms. First, SET7-mediated H3K4me1 activates transcription of spliceosome components, promoting inflammatory and oxidative responses. Second, SET7 directly mono-methylates endothelial nitric oxide synthase (eNOS) at lysine 494, disrupting calmodulin binding, hence nitric oxide production. Importantly, activation of SET7-splicing axis and the disruption of eNOS-calmodulin interaction were also observed in arteries from patients with established atherosclerosis and hyperglycemia, supporting their clinical relevance. Overall, this study provides a rationale for targeting SET7 as a putative therapeutic strategy to reduce the burden of vascular complications in patients at high cardiometabolic risk.

